# Individual symbiont strains can compensate for complex microbiome loss in the bull-headed dung beetle, *Onthophagus taurus*

**DOI:** 10.64898/2026.11.19.767994

**Authors:** Joshua A. Jones, Irene L. G. Newton, Armin P. Moczek

**Affiliations:** Department of Entomology, University of Minnesota, St. Paul, Minnesota, USA; Department of Biology, Indiana University, 915 East 3rd Street, Bloomington, IN 47405, USA

**Keywords:** host-microbe interactions, nutritional symbiosis, vertical transmission, Scarabaeoidea, Coleoptera, coprophagy, *Acinetobacter*, *Brucella*, *Pseudomomas*, *Dysgonomonas*

## Abstract

Many animals and plants have evolved to rely on microbiome mediated interactions with their environment. Across diverse insect groups, many are incapable of either breaking down complex carbohydrates or synthesizing essential nutrients despite consuming diets which should require one, or both, functions. *Onthophagus* beetles are one such host, with recent work suggesting their ability to digest their dung diet, typically rich in tough to digest polysaccharides and poor in nutrients, is likely bolstered by heritable symbionts. Despite there being over 2,000 species of *Onthophagus*, and over 5,000 species of “true” dung beetles (Scarabaeinae), little is known about how the microbiomes of these beetles modulate their interactions with their dung diet. The work presented here assesses the possible significance of four individual microbiome members, present during *Onthophagus taurus* development. Results show that three of the four isolates tested partially recovered normative developmental outcomes in *O. taurus.* Specifically, inoculation with a strain of either *Acinetobacter*, *Brucella*, or *Pseudomonas* increased pupal size relative to beetles deprived of the maternal inoculate. This mirrors the effects of individuals receiving the full natural inoculate, which contains hundreds of bacterial species. Further, using whole genome sequencing, we form a hypothesis of what nutrients these isolates provide to benefit *O. taurus* development. The findings presented here further highlight *Onthophagus* dung beetles as powerful systems to study the complex ways in which microbiome function can interact with diet to influence developmental and evolutionary outcomes.

**Importance:** Microbiotas are communities of microbes that inhabit many animals and plants and are often major factors in their hosts’ health, development, and evolution. There are, however, relatively few systems where work has gone beyond finding associations between hosts and microbes and into testing microbiome reliance or identification of microbiome functions. This lack of causal evidence has limited our ability to understand the factors controlling host-symbiont interactions, how these pairings are formed, and how these relationships change throughout evolutionary time. This work expands our understanding of host-microbe interactions in, *Onthophagus* dung beetles, an understudied system which has considerable ecological and species diversity. By validating the relevance of the microbiome to dung beetle development and showing that the microbiome can be experimentally manipulated, we hope to highlight that experimentation in dung beetles may serve to further our understanding of the evolution and maintenance of host-symbiont interactions.

## Introduction

Animals demonstrate a wide range of complex traits which improve their ability to benefit from their dietary resources. Much research into these adaptations has focused on host traits but the microbiome, communities of microbes inhabiting a host, has emerged as a key extragenetic effect influencing resource use. For example, the microbiome is key to the dietary ecology of many insects, allowing for the digestion of complex polysaccharides(1–5), the biosynthesis of key nutrients(6–9), and the detoxification of dietary toxins(10,11) required for those insects to subsist on their main diet sources. Thus, the interactions between host and microbiome can have considerable influence on the fitness and evolution of many insects. To ensure that these interactions are maintained, some insects have evolved mechanisms to retain individual microbes or entire microbiota across life stages and generations(12–15). Despite the likely importance of microbiomes to many insects, the specific metabolic function, developmental effects, and mechanisms of maintenance are well understood in only a select few model organisms(16–19).

*Onthophagus* beetles represent a useful system to broaden our understanding of host-microbiome interactions. This genus of dung beetle is highly speciose, containing an estimated 2,500 species, the majority of which are entirely reliant on dung as a diet and reproductive resource (20,21). Many of these coprophages are specialized on herbivore dung, a recalcitrant resource high in tough to digest polysaccharides and low in essential nutrients(22,23). Adult beetles are filter feeders which consume the nitrogen rich liquid portion of the dung, while larvae, sequestered in underground *brood balls* of dung provisioned by parents, consume the less compositionally understood fibrous portion(23). This fibrous portion is likely a difficult diet for these beetles as it contains mainly plant cell wall materials that most animals(24,25), including dung beetles(26), lack the enzymes to effectively digest. Like many other insects specialized on recalcitrant diets, dung beetles therefore rely on a heritable microbial community to overcome the challenge posed by their diet. *Onthophagus* beetles, such as *O. vacca* and *O. medius*(27), and an outgroup of *Onthophagus*, *Digitonthophagus gazella*(28,29), suffer significant developmental deficits when deprived of the *pedestal*, a fecal swab left by mothers which transfers the microbiome across generations(13,28). Recent work has begun to document the microbial community of the pedestal and has identified several common bacterial taxa, derived in part from the food the mother consumes, such as *Acinetobacter* and *Pseudomonas*, but also taxa absent from the diet which nevertheless dominate the microbiome at different points in development, such as *Dysgonomonas* and *Brucella*(26,30). In this study we sought to further extend our understanding of the importance of maternally inherited microbiota to dung beetle development, and the mechanisms by which their benefits may be conferred. Specifically, we sought to investigate whether *Onthophagus taurus* - a common model dung beetle in evolutionary developmental biology thus far understudied with respect to its microbiome biology - also relies on the pedestal microbiota to support normative development. We then aimed to determine whether isolates from four bacterial genera that were previously found to dominate dung beetle microbiota *(Acinetobacter, Pseudomonas*, *Dysgonomonas, Brucella*) are by themselves sufficient to support normative *O. taurus* development. Lastly, we sought to characterize what specific metabolic resources these isolates may be providing to benefit host development.

To do so, we contrasted the developmental performance of hosts with access to the full complement of pedestal derived microbiota and those deprived of these, but provided one of four bacterial inoculants, to those deprived of their pedestal and not provided any inoculant. Our results suggest that *O. taurus*, similar to other tested dung beetles, grow to larger sizes when provided the maternal pedestal, and that pedestal-deprived larvae provided with either *Acinetobacter*, *Brucella*, or *Pseudomonas*, but not *Dysgonomonas*, develop into heavier pupae than pedestal deprived larvae. Additionally, we found that this rescue translates into adult size in *Brucella* and *Pseudomonas* and that *Dysgonomonas* alone impacts development time. Lastly, we assembled the genomes of the four focal microbial taxa and documented that all except *Dysgonomonas* encode the complete modules for the biosynthesis of lysine and biotin, both of which are important for arthropod development(31–33), are not encoded within the dung beetle genome(26), and are unlikely to be abundant in their diet(22,23). Taken together, this work establishes a model and framework for future exploration into the interactions between host diet and microbiome function, which may help shed light onto the evolution of complex symbioses such as these.

## Materials and Methods

### General beetle husbandry

*Onthophagus taurus* adults were collected near Chapel Hill, North Carolina, U.S.A., in early summer 2024 and maintained in colonies of F0 individuals. Colonies were maintained at 24°C, 40% humidity, with a 16:8h light:dark cycle and fed twice a week with homogenized dung collected from Marble Hill Farm in Bloomington, IN.

### Microbe isolation, storage, and identification

Bacterial isolates used in this experiment were harvested either from *O. taurus* directly (*Acinetobacter*, *Brucella*, and *Dysgonomonas*) or from the maternally provisioned pedestal (*Pseudomonas*). All beetles dissected for plating were produced by transferring 6 adult females and 3 males into breeding containers as detailed in Shafiei et al. (2001)(34) and providing them with *ad libitum* dung for 7 days to allow for the production of brood balls containing eggs. Pedestals were collected directly from brood balls while larvae, pupae, and adults were collected by transferring eggs to 12-well plates, with individual wells provisioned with homogenized dung from cows previously feeding on grass, and allowing eggs to hatch and to progress through development within the well until they reached the desired life stage (as detailed in Shafiei et al., 2001(34)). All individuals were anesthetized at-20°C for 10 minutes and surface sterilized with a sterilization solution (double-distilled water with 1% bleach and 0.1% triton-X 100) prior to dissection, homogenization, and plating. An *Acinetobacter* isolate was obtained from the adult mid and hindgut plated onto a custom media recipe (1x M9 Salts, 1mM MgSO_4_, 0.1mM CaCl_2_, 0.01mM cellulose, and 50mg each of cysteine, glycine, proline, serine, threonine, phenylalanine, and valine, in 1 L double-distilled water) in normal atmospheric conditions at 24°C. *Brucella* was isolated from a whole pupal homogenate on *Brucella* agar (CRITERION, Cat. no. C5300) in normal atmospheric conditions at 24°C. *Dysgonomonas* was isolated from the larval hindgut on Bacteroides Bile Esculin agar (S2 Media, Cat. no. CHM01P625) in a BD GasPak EZ incubation container (REF 260672) using GasPak EZ Anaerobe Container System Sachets (REF 260678) at 24°C. Finally, *Pseudomonas* was isolated from pedestals collected from unhatched eggs and plated on *Pseudomonas* isolation agar (NEOGEN, Cat. no. NCM0150A) in normal atmospheric conditions at 24°C. All selected colonies underwent two rounds of streaking for isolation before being grown up in LB broth (BD Difco, Cat. no. DF0446-17-3) to produce adequate biomass for both PCR for identification and storage in a 50:50 ratio of sterile 50% glycerol and culture broth for storage in a-80°C freezer. Isolates were initially identified by sequencing the 16S rRNA gene with 27F and 1492R primers (27F: AGA GTT TGA TCM TGG CTC AG; 1492R: GGT TAC CTT GTT ACG ACT T; M represents A or C) and BLASTing(35) the resulting sequence against NCBI’s nr database(36).

### Beetle rearing for inoculation experiments

Beetles used in this experiment were reared by setting up breeding containers, as described above, allowing F0 beetles to produce brood balls for three days. The shorter reproduction window helped to ensure that brood balls still contained eggs instead of hatched larvae. After harvesting brood balls, eggs were randomly assigned to a treatment group, processed as detailed below, and placed into a 12-well plate packed with sterilized dung (as in Schwab et al., 2016(28)) obtained from hay-fed cows. Hay derived dung was used as it represents a more difficult diet compared to dung produced by grass-fed cows (Rohner & Moczek 2021(37)), thus possibly increasing reliance on metabolic services provided by the microbiome. Treatment groups included: M+ (individuals developing in the presence of their pedestal and egg surface microbiomes), M-(individuals deprived of their pedestal and egg surface microbiomes), and inoculant groups (individuals deprived of their pedestals and surface microbes but supplemented with a single inoculant of either *Acinetobacter*, *Brucella*, *Dysgonomonas*, or *Pseudomonas*). Eggs assigned to M+ were removed from their brood ball and placed directly into their assigned well along with their pedestal. M-individuals were removed from their brood ball and sterilized (rinsed twice with 200μL of sterilization solution, described above, and two more times with 200μL of sterile PBS) before being placed into their assigned well. Importantly, past work suggests that this method does not result in axenic beetles(38) yet it effects the microbiome enough to have clear developmental effects on the host(28). Inoculant eggs were treated like M-eggs, plus provided an addition of 100μL of their assigned inoculant, suspended in PBS, instead of sterile PBS. Inoculants were prepared by adding ∼5μL glycerol stock to 5mL LB broth and continual rotating until experiment setup (1-2 days). Prior to inoculation, the inoculant was washed by centrifuging at 16,100rcf for 1 minute followed by removal of the supernatant, resuspension in 1mL sterile PBS, centrifugation for another minute, removal of the supernatant, and resuspension in 1mL sterile PBS. The bacterial load of this suspension was measured using a hemocytometer before being diluted to approximately 10,000 bacteria per 100μL. Finally, 100μL of sterile PBS was doused on all M+ and M-eggs, to eliminate any potential effects of starting moisture. This protocol was repeated across 5 batches, at which point at least 216 eggs were processed allowing for 36 eggs per group.

### Developmental metrics

To assess the effect of different microbiome treatments on *O. taurus* development, we collected data on hatch date, larval weight, pupation date, pupal mass, sex, male horn morphology and length, eclosure date, survival throughout development, female fecundity, female adult survival, and adult body size. All beetle handling throughout development was conducted with sterilized forceps, which were re-sterilized in bleach solution followed by a sterile PBS wash between handling different individuals. Hatch date, pupation date, eclosure date, and survival throughout development was determined by checking each well daily. During this checking, we took care to cause minimal disruption to the larval brood ball environment to minimize possible negative effects on subsequent development(29,39). Larval and pupal mass was determined by weighing individuals using an analytical balance (Mettler Toledo, AL54), individuals were all weighed on fresh weighing paper (VWR North America, Cat. no. 12578-121) to prevent cross contamination. Larval weight was recorded on day seven after hatching and pupal weight was taken on day 2 after pupation. Sex and male horn morphology was determined through visual inspection of pupal head horns, both of which were verified upon eclosure by inspecting adult horn morphology. After eclosure, adult beetles were put into inoculant-specific plastic containers with moist soil and dung to finish sexual maturation and allow for fecundity measurements. Female beetles were marked on their elytra (following Silva & Hernández 2015(40)) so they could be identified and moved to a breeding container after three weeks of maturation. Only a single female produced brood balls in the fecundity assay, so fecundity across groups could not be analyzed. After females finished the fecundity assay, they were preserved in centrifuge tubes of 70% ethanol. Males were maintained in their inoculation-specific containers, to assist with sexual maturation, until the final females began the fecundity assay, at which point the remaining males were also preserved in 70% ethanol. Beetles found dead throughout this time window were also preserved in 70% ethanol. Some individuals rapidly deteriorated before they could be collected, leading to a lower sample size for adult size. After all remaining adults were preserved, thorax width and horn length was measured by taking photos using a mounted camera (Pixelink PL-D7912CU-T) on a stereo microscope (Leica MZ16). A photo of a micrometer (Graticules PYSER-SGI LTD. 02A00400) was taken with each batch of photos to provide a standardized ruler for calibration. Measurements were obtained using Fiji (Schindelin et al., 2012(41)). Thorax width was obtained by measuring the width of the thorax (as in Moczek & Emlen 1999(42)) and horn length was measured by averaging left and right horn length measured from the edge of the eye cavity to the tips of the horns (as in Moczek & Emlen 1999(42)).

### Statistical analysis

All statistical analysis was conducted using R version 4.2.2(43). The effect of treatment on larval mass, pupal mass, adult thorax size, and developmental rate relative to the microbiota diminished group was analyzed using trt.vs.ctrl method in *emmeans*(*44*), which implements Dunnett’s many-to-one comparison method(45) on a linear mixed model, made using *lmer*(*46*). This linear model contained the focal trait as the dependent variable, microbiome as the sole independent variable, and batch, plate, and well as random effects, except in the model for adult size where these random effects were not included. Similarly, the effect on survival and horn morphology was analyzed using *emmeans* to implement a Dunnett’s test to analyize a generalized linear model(47–50), made using *glmer*(*46*), with the focal variable as the dependent variable, microbiome as the independent variable, and batch, plate, and well as independent variables. Dunnett’s test was used to avoid the loss of power that would result from running an ANOVA testing all comparisons instead of those between the treatments and the pedestal deprived beetles. Data used in these analyses are included in the data repository. Data wrangling and visualization was completed using *cowplot* v1.2.0(51), *emmeans* 2.0.4(44), *ggdist* v3.3.3(52), *ggtext* v0.2.0(53), *lmerTest* 3.2.1(54), *tidyverse* v2.0.0(55), list made using *grateful* v0.3.1(56).

### Genome sequencing, assembly, and analysis

To investigate which metabolic functions may account for the microbial influences on development, we assembled genomes for the four isolates used in this study. Isolates were grown in the lab and DNA extractions, barcoding, and sequencing were carried out at SeqCoast Genomics (Portsmouth, NH, USA) for both Illumina short read and Oxford Nanopore long read sequencing. *Brucella* was grown and sent for sequencing in Fall 2023 while the remaining isolates were sent in Winter 2024 resulting in minor differences in processing and software. Differences in Winter 2024 sample processing are noted in parentheses following the version used to process *Brucella* in Fall 2023. Samples were lysed with MAGMAX Microbiome bead beating tubes before DNA extraction using Qiagen DNeasy 96 PowerSoil Pro QIAcub HT Kit #47021. Illumina reads were prepared using an Illumina DNA prep tagmentation kit and unique dual indexes. Sequencing was performed on the Illumina NextSeq2000 platform using a 300-cycle flow cell kit to produce 2×150bp paired reads. 1-2% PhiX control was spiked into the run to support optimal base calling. Read demultiplexing, read trimming, and run analytics were performed using DRAGEN v3.10.12 (v4.2.7) on the NextSeq2000. Nanopore files were prepared using the Oxford Nanopore Technologies SQK-NBD114 native barcoding kit. Long Fragment Buffer was used to promote longer read lengths. Sequencing was performed on the GridION platform (PromethION 2) using a FLOW-MIN114 Spot-ON Flow Cell (FLO-PRO114M), vR10 with a translocation speed of 400bps. Base calling was performed on the sequencing platform using the super-accurate basecalling model, MinKNOW v23.07.5 (v24.06.10) and Guppy v7.0.9 (Dorado v7.4.12).

All isolate genomes followed a unified workflow for quality-controlling, assembly, and annotation. Illumina reads were trimmed using Trimmomatic v0.39 (LEADING:3 TRAILING:3 SLIDINGWINDOW:4:20 MINLEN:36)(57) followed by checking with FastQC v0.12.1(58). Nanopore reads shorter than 1,000 base pairs (bps) or a phred score lower than 20 were removed using seqkit v2.8.2(59,60). Unicycler v0.5.0 was used to assemble the genomes using both Illumina and Nanopore reads(61). Assembly completeness and contamination statistics were assessed using CheckM v1.2.3(62). Subsequent identification and annotation work, unless otherwise noted, was conducted in KBase(63). Taxonomic identification was finalized using GTDB-tk v2.3.2(64–71) to find the closest genetic relative in GTDB database release 214. RNAs and genes were predicted, and annotated, using RAST-tk v1.073(72–74) and Prokka(75) within KBase, and the NCBI Prokaryotic Genome Annotation Pipeline(62,76–84) outside KBase. RASTtk output is available in the RASTtk.GFF files included in the data repository. Using the RAST-tk output, the *Dysgonomonas* and *Pseudomonas* genome sequences were rotated to begin at *dnaA*. This was done here because automatic rotation by Unicycler was unsuccessful for these genomes. These two genomes were then reannotated so that gene locations aligned with the rotated sequences. Each genome assembly is available as BioProject PRJNA1363681. Protein sequences were assigned Carbohydrate-active enzymes (CAZy) functions based on alignments to the dbCAN2 HMM collection using Search with dbCAN2 HMMs of CAZy families which uses HMMER v3.3.2 and the dbCAN2 HMM collection(64,70,85). Abundance of CAZy classes were analyzed for each isolate genome using program outputs in R version 4.2.2(43). Finally, biosynthesis potential was predicted by determining the proportion of enzymes in KEGG modules(86) with orthologs in the respective isolate genome, using the RASTtk annotations, which was compared to genes predicted for the *O. taurus* genome(87). Listings of KEGG enzyme nomenclature numbers along with which genomes contained orthologous genes are available in the data repository.

## Results

This study sought to investigate the degree to which specific microbial community members may be sufficient to support normative development in the bull-headed dung beetle *O. taurus* in place of the normally complex, maternally inherited microbiome, and to characterize the specific metabolic benefits these individual isolates may be providing to their hosts. To accomplish this, we measured the developmental outcomes of *O. taurus* individuals inoculated with one of four bacterial isolates, selected based on (a) their presence throughout important phases of host development or (b) their potential to harbor key metabolic pathways, or both. Developmental outcomes of inoculated beetles were compared against individuals who developed either in the presence of, or absence from, their pedestal and egg surface microbes, i.e. the hosts’ main natural sources of microbial acquisition. This effort was then paralleled by a detailed genomic analysis of microbial isolates. We show that the benefits provided to *O. taurus* by their pedestal microbiota can be partially recovered with the presence of individual microbial community members, and identify putative metabolic benefits symbionts may provide which could lead to the enhanced developmental outcomes we observed.

### Maternal pedestals influence pupal mass and adult size

To measure the effect of pedestal microbes on development, we assessed differences between individuals who developed in the presence of their pedestal and those who were surface sterilized as eggs and deprived of their pedestals. Using a Dunnett’s test, we observed that the presence of the pedestal increased pupal mass and adult size, but not other metrics, relative to the microbiome depleted controls. Specifically, individuals with their pedestals were an estimated 17.17 mg heavier (t = 3.37, p = 0.001, Figure 1A) and 0.28 mm larger on average (t = 2.866, p = 0.048^-1^, Figure 1C). However, there were no significant differences in the larval mass (23 mg, t = 0.493, p = 0.623, Supplemental Figure 1A), time to pupation (-0.26 days, t =-0.342, p = 0.733, Figure 1B), time to eclosure (-0.305 days, t =-0.368, p = 0.713, Supplemental Figure 1B), likelihood of survival to eclosure (0.751 odds ratio, z =-0.500, p = 0.617, Supplemental Figure 2A), or likelihood of males developing a major horn morphology (0.281 odds ratio, z =-1.513, p = 0.130, Supplemental Figure 2B) between the treatments. These results support earlier studies that document the positive impact of the maternally inherited microbiome on beetle development(27–29) and thereby validate the experimental approach taken in this study.

**Figure 1.**
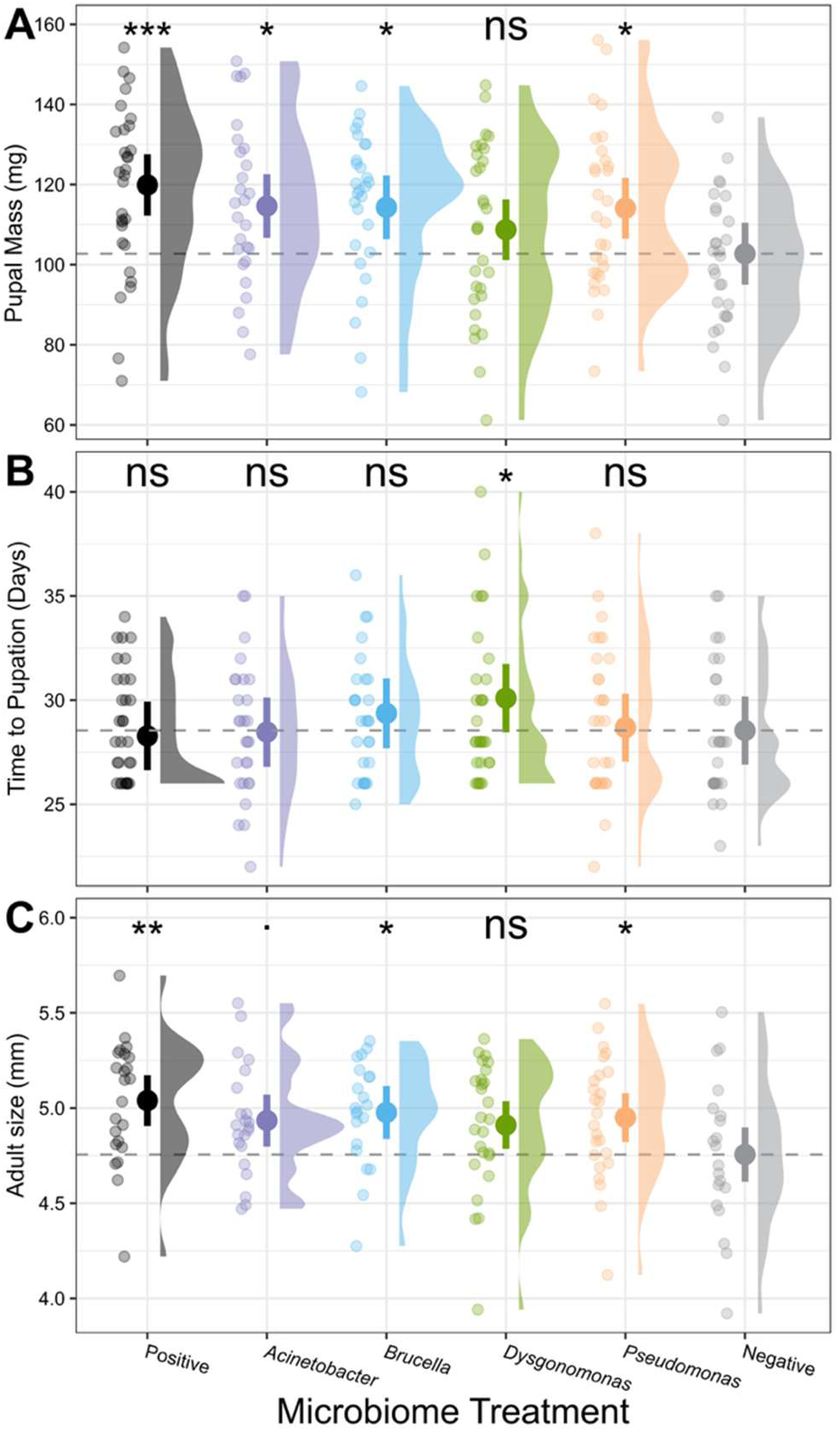
Partial removal of the maternally inherited microbiome, via pedestal removal and egg surface sterilization, negatively influences pupal mass and adult size but inoculation with select microbes can provide partial rescue. A) Pupal mass, B) time to pupation, and C) adult size across treatments. Large points represent estimated marginal means within each metric and treatment group, error bars are 95% confidence intervals, and small points are individual data points. Dashed line is a reference to the mean of the microbiome diminished treatment. Asterisks represent significant difference from the microbiome diminished treatment (ns > 0.1,. <= 0.1, * <= 0.05. ** <= 0.01, *** <= 0.001)

### Select isolates partially recover development

To determine if single inoculants were sufficient to recover the developmental benefit of the pedestal, the developmental outcomes of beetles receiving one of four bacterial inoculants were also compared to the microbiome depleted controls. From the same Dunnett’s test reported above, we saw that inoculation increased the average pupal mass in *Acinetobacter* (+11.90 mg, t = 2.312, p = 0.022, Figure 1A), *Brucella* (+11.60 mg, t = 2.273, p = 0.024), and *Pseudomonas* (+11.36 mg, t = 2.249, p = 0.026) but not *Dysgonomonas* (+6.01 mg, t = 1.208, p = 0.229). This mass increase only translated into an adult size increase in *Brucella* (+0.221 mm, t = 2.184, p = 0.031, Figure 1C) and *Pseudomonas* (+0.194 mm, t = 2.000, p = 0.0476), with *Acinetobacter* (+0.178 mm, t = 1.786, p = 0.765) and *Dysgonomonas* (+0.155 mm, t = 1.609, p = 0.110 reaching adult sizes larger but insignificantly different than the controls. The only treatment, including the pedestal treatment, differing from the control on any other metric was *Dysgonomonas* which took significantly longer to reach pupation than the controls (+1.555 days, t = 2.115, p = 0.036, Figure 1B). Supplemental Table 1 contains the outputs from the Dunnett’s tests, no results except those reported here were statistically significant.

### Isolate genome statistics

We then assembled the genomes for each of the isolates used in our treatments in hopes of identifying which specific metabolic functions may underlie the effects we observed on host development. Initial sequencing yielded a total of 4.9 billion sequenced base pairs, 1.7 from Illumina and 3.2 from Nanopore sequencing, respectively. Following our quality-control steps, assembly with Unicycler produced circular genomes for each isolate with only the *Pseudomonas* genome not being contiguous. CheckM found that all four genomes contain all ubiquitous, single copy genes expected in their genomes, suggesting nearly 100% completeness. PGAP predicted between 5,089 (*Pseudomonas*) and 3,629 (*Dysgonomonas*) genes each, including 4,989 of which fit into 359 different SEED functional roles (Supplemental Figure 3). Further summary statistics are included in Table 1.

**Table 1.** Summary statistics for isolate genomes used in inoculation study and functional analyses.

| Isolate | Acinetobacter | Brucella | Dysgonomonas | Pseudomonas |
|---|---|---|---|---|
| BioSample | SAMN53215226 | SAMN53215227 | SAMN53215228 | SAMN53215229 |
| Contigs | 1 | 2 | 1 | 8 |
| Scaffolds | 1 | 2 | 1 | 1 |
| Genome Size (bp) | 4,453,094 | 4,732,911 | 4,457,789 | 5,697,305 |
| Lengths | 4,453,094 | 2,701,023<br>2,031,888 | 4,457,789 | 5,697,305 |
| GC% | 37.62% | 56.04% | 62.45% | 36.49% |
| Genus Identification | *Acinetobacter* | *Brucella* | *Dysgonomonas* | *Pseudomonas* |
| Closest ANI | GCF_902825285.1 | GCF_000017405.1 | GCF_011299555.1 | GCF_000935045.1 |
| Closest Relative Taxonomy | Acinetobacter sp. Marseille-Q1620 | Brucella anthropi ATCC 49188 | Dysgonomonas sp. HDW5A | Pseudomonas sp. 10-1B |
| Percent Alignment to Closest Relative | 98.64% | 97.56% | 96.47% | 97.53% |
| Closest Placement Alignment Fraction | 0.96 | 0.89 | 0.91 | 0.84 |
| Raw Illumina reads | 2,953,796 | 2,227,942 | 3,568,772 | 3,108,796 |
| After QC | 2,850,110 | 2,141,226 | 3,454,624 | 2,981,892 |
| Raw Illumina bps | 429,332,998 | 332,443,107 | 517,679,262 | 451,512,551 |
| After QC | 407,254,563 | 308,473,984 | 493,992,799 | 424,895,735 |
| Raw Nanopore reads | 71,616 | 65,943 | 298,801 | 49,230 |
| After QC | 43,726 | 29,774 | 88,842 | 27,698 |
| Raw Nanopore bps | 968,269,725 | 649,137,201 | 879,569,207 | 699,613,377 |
| After QC | 636,794,581 | 328,921,656 | 464,561,137 | 429,756,703 |
| CheckM Completeness | 100.00% | 100.00% | 100.00% | 100.00% |
| CheckM Contamination | 0.03% | 0.00% | 0.00% | 0.30% |
| RASTtk Gene Count | 7,010 | 8,694 | 5,756 | 10,131 |
| Prokka Gene Count | 4,248 | 4,484 | 3,751 | 5,133 |
| PGAP Gene Count | 4,120 | 4,350 | 3,629 | 5,089 |

### *Isolate genomes differ in* CAZy abundance and nutrient biosynthesis potential

To obtain a general estimate of polysaccharide utilization potential across the isolates, we quantified the abundance of genes aligning to the six CAZy enzyme classes. We found a total of 453 genes aligning to the CAZy database (Figure 2). The majority of these (265) are in the *Dysgonomonas* genome, followed by *Brucella* and *Pseudomonas* at 69 each, with *Acinetobacter* having the least at 49. Next, we determined the potential for each isolate to synthesize the ten essential amino acids which are absent from the *O. taurus* genome(26). Analysis of these pathways revealed none of these isolates contained complete modules for all ten essential amino acids (Figure 3). *Brucella* had the most complete modules with 9/10, followed by *Pseudomonas* (8/10), then *Acinetobacter* and *Dysgonomonas* with the least (7/10). Finally, we predicted if these isolates could synthesize several key B vitamins. Here, we found that *Pseudomonas* has the highest potential (4/7), followed by *Acinetobacter* and *Dysgonomonas* (3/7), with *Brucella* having the least (1/7). Of note, both lysin and biotin pathways were complete in all but the *Dysgonomonas* genome, raising the possibility that these specific nutrients may play key roles in the developmental outcomes we observed.

**Figure 2.**
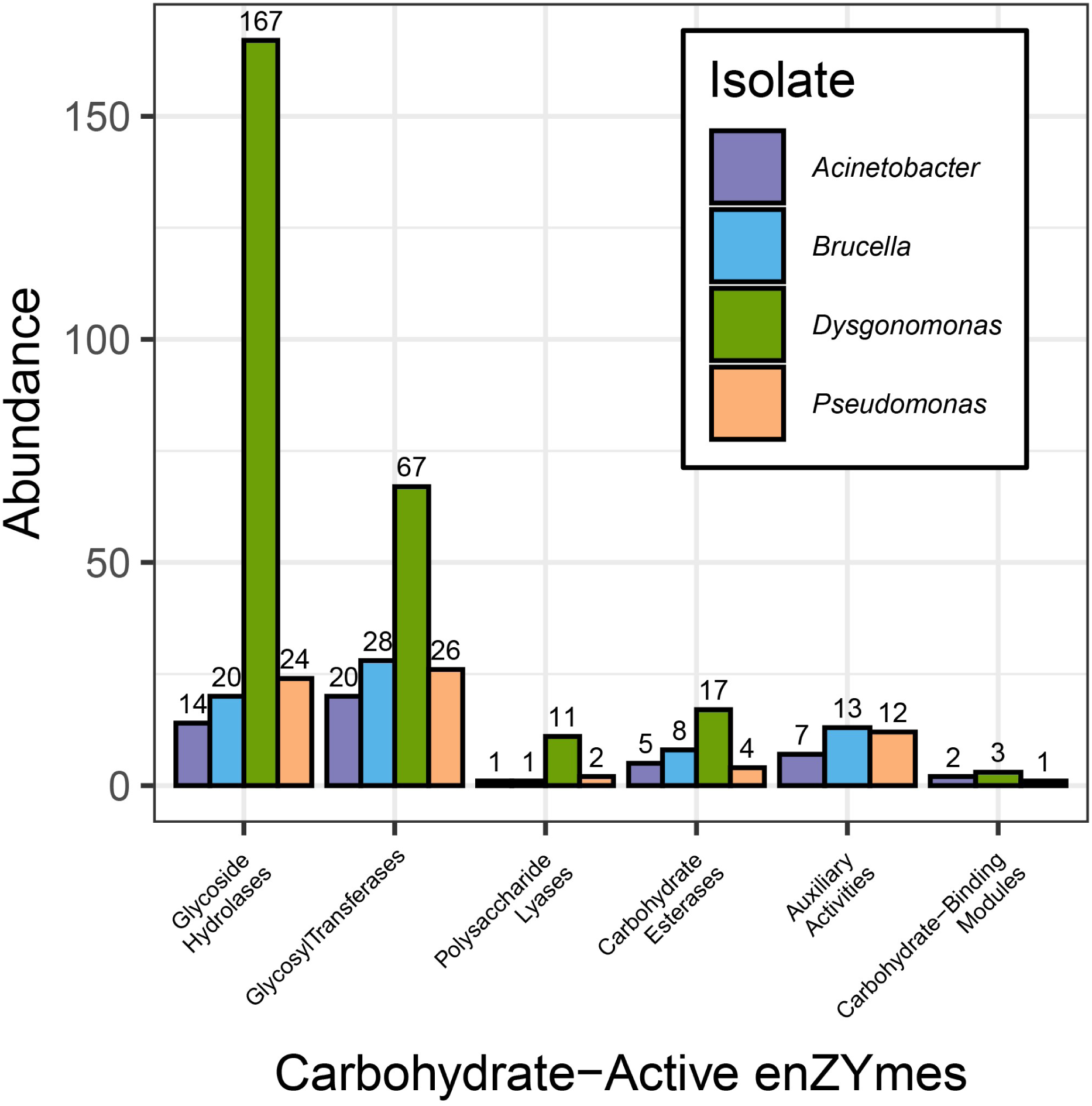
Abundance of enzymes assigned to CAZy categories across isolate genomes. Bar colors differentiate isolate genomes and numbers above bars are counts of total enzymes.

**Figure 3.**
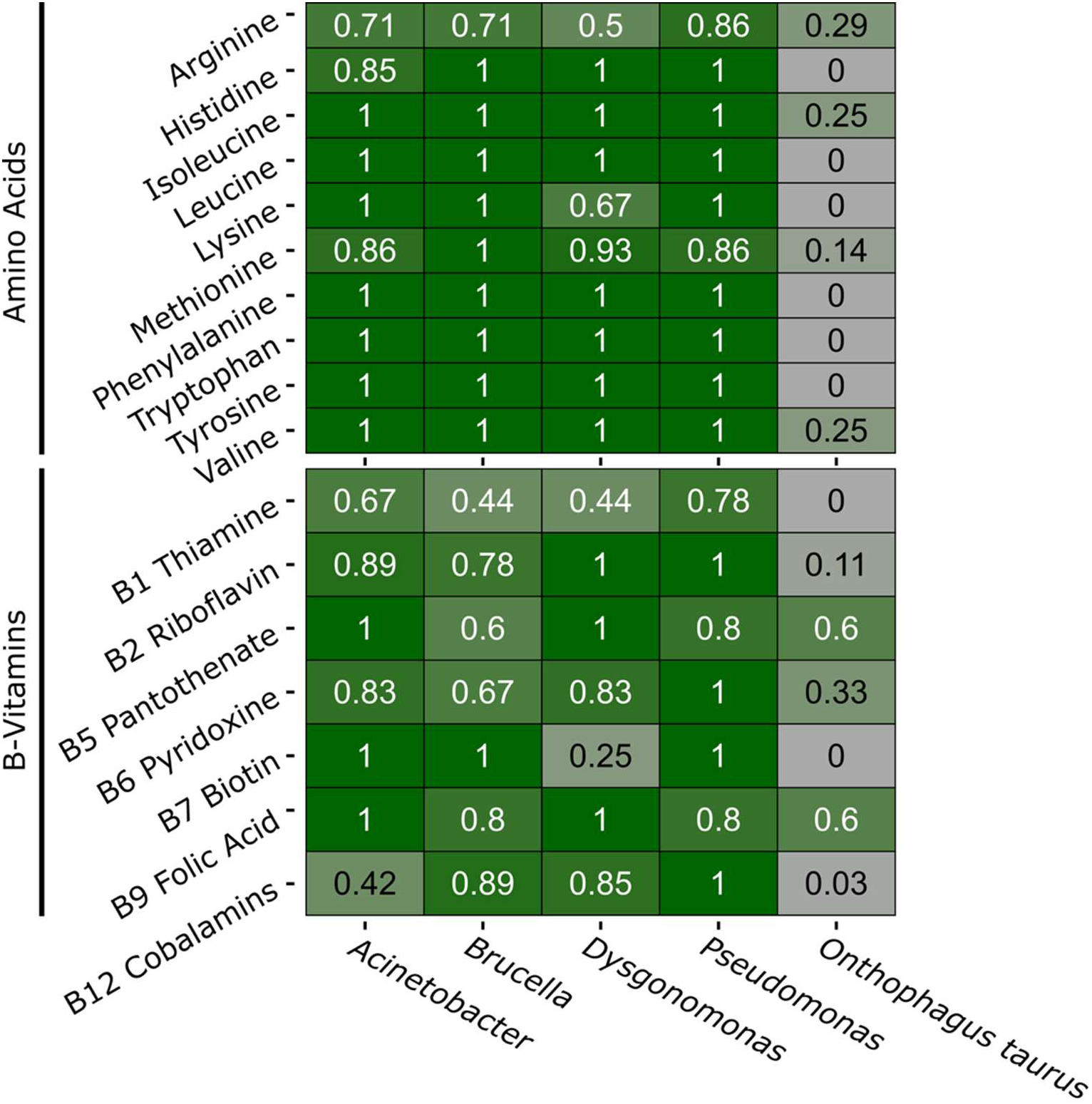
Completeness of nutrient pathways across isolate and host genomes. Color scale shows what proportion of genes within the KEGG module have orthologs within the genome.

## Discussion

Many multicellular organisms rely on communities of microbial organisms for major components of their ecology, such as resource digestion(1–5) and nutrient biosynthesis(6–9), resistance to toxins and pathogens(10,11,88), or to support growth and development(89,90). These communities, thus, represent major factors influencing how organisms respond to their environment, allowing for the possibility that differences in community structure may partially account for differences in developmental and ecological outcomes(91). Here, we use a dung beetle, *Onthophagus taurus*, to determine whether exposure to specific microbial inoculants – selected based on their abundance and metabolic potential - are partially or fully sufficient to provide the developmental benefits conveyed by the presence of the microbial community as a whole. We then assembled and annotated the genomes of our focal microbes to deepen our understanding of the possible metabolic functions underlying the observed developmental outcomes.

### Individual isolates are capable of partially recovering the developmental effect of the pedestal microbiome

Our developmental assay demonstrated that, similar to other tested dung beetles(27–29), the maternally derived pedestal had a significant effect on pupal mass and adult body size in *Onthophagus taurus*. Further testing revealed that the developmental benefit of the pedestal can be partially recovered by microbial taxa commonly associated with, and cultured from, *O. taurus*. Specifically, inoculating pedestal-deprived larvae with an isolate of *Acinetobacter*, *Brucella*, or *Pseudomonas* resulted in pupal masses higher than those of the uninoculated controls. Further, these effects carried over to the adult stage in the *Brucella* and *Pseudomonas* groups, with a trend towards larger size in the *Acinetobacter* group as well. Though this design was not sufficient to measure differences in fitness across treatments, this difference in size may serve as a selective force maintaining beetle-microbe interactions over preventing microbiota inheritance. Further, since we did not measure microbiota composition throughout development, we cannot conclude whether these results stem from direct interactions with the inoculated isolates or the indirect effects of those inoculations. While the beneficial influence of the pedestal on development has been reported previously in other dung beetle species (*Digitonthophagus gazella*(28,29) & *Onthophagus vacca* & *O. medius*(27)) this is the first evidence documenting such an effect in *O. taurus*. Additionally, this is the first study demonstrating that the effects of individual isolates can be measured in dung beetles, priming these insects for future work to provide causal evidence linking isolates to host responses.

Lastly, previous work on dung beetle-microbiome interactions also demonstrated that the microbiome may influence developmental timing (i.e., time to pupation or adulthood(27–29,92)). However, in this study the timing of key developmental events was similar across all our treatments (similar to(93)) aside from the *Dysgonomonas* group, which experienced a slowed development to pupation. This discrepancy could be due to either a loss of some influential symbionts during lab rearing or population/species level differences in the microbiome’s influence on host traits, both of which have been demonstrated in various studies across other taxa(94–98). Further work to understand the ecological and evolutionary context influencing which traits symbionts impact may be of interest.

### Biosynthesis of key vitamins and amino acids may play a major role in microbe assisted growth

Intriguingly, inoculation with *Dysgonomonas* was the only experimental manipulation we executed that failed to yield an effect on pupal mass, even though *Dysgonomonas* constitutes the most dominant taxon within the *O. taurus* larval microbiome(30). While abundance can be an indication that a symbiont is functionally relevant, it is obviously not a perfect signal. Similarly, it is also possible that our experimental design removed some factor needed for *Dysgonomonas*’ functional role to be apparent. Further study of the functional diversity within symbionts and the resource diversity within the diet are thus needed to uncover the factors influencing *Dysgonomonas*’ abundance throughout development.

Alternatively, *Dysgonomonas* may lack critical biochemical abilities compared to other isolates that might explain why, when inoculated in isolation, it failed to positively affect development. To begin addressing this question we utilized our genome assemblies to compare the metabolic potentials across the four isolates’ genomes to identify putative metabolic functions that may influence beetle growth, focusing on essential nutrients biosynthesis in general and amino acid synthesis specifically(22,26,99) (but also see Holter 2016(23) who discusses that amino acids may be plentiful in the dung microbiome). Our results suggest that the genomes of *Acinetobacter*, *Brucella*, and *Pseudomonas*, all of which increased pupal mass and had variable effects on adult size, contained the biosynthetic pathways required to synthesize seven of the ten essential amino acids. Of these, lysine may be of particular interest: symbionts provide lysine in other insects that utilize amino acid deficient diets (honeybees(100) & whiteflies(101)), yet the necessary pathway was absent in *Dysgonomonas* but not the other three isolates used in this study. Further, we identified only a single B-vitamin pathway shared across the isolates that improved development - biotin – which, similar to the lysine biosynthesis pathway, was missing from the *Dysgonomonas* genome. Several studies on other insects have assessed the effect of biotin deprivation and documented comparable developmental outcomes such as limited or arrested development in bedbugs(31) and reduced mitosis rates in the cells of fruit flies(33).

Again, as we did not validate the continued presence of our inoculants in the microbiome, nor did we employ further-omics to detect changes in these nutrients or symbiont enzymes, the implication of these pathways in our results is entirely theoretical. Future work manipulating the abundance of these putatively important nutrients alongside candidate symbionts, or quantifying the effects of symbionts on nutrient abundance, will help determine the function of these symbionts and expand our knowledge of the nutritional physiology of these insects.

## Conclusion

In this study we provide further evidence for the developmental benefits of individual bacterial isolates and suggests for the first time specific microbial taxa and potential microbial byproducts of importance to dung beetle hosts. Future studies may manipulate the presence of these, or other, isolates in dung beetles, along with supplementation of nutrients into the dung diet, to further investigate their significance to *O. taurus* and other members of this speciose genus. Beyond this, the work presented here sets up dung beetles as a potentially powerful study system to help disentangle the complex relationships between host, microbiome, and resource ecology. Specifically, *O. taurus* now represents a useful model in which bacteria with defined metabolic potentials can be added or removed, in concert with manipulations of the environment, a capability which few non-traditional model systems have achieved.

## Dava availability

Data used to produce developmental metrics, RASTtk outputs from each isolate, and binary listing of the presence/absence of orthologous genes for nutrient biosynthesis are included in the data repository on figshare (10.6084/m9.figshare.33427252). Isolate genomes are included in BioProject PRJNA1363681 and BioSamples SAMN53215226-SAMN53215229.

## Acknowledgements

We would like to thank Tracey Rettig for collecting *O. taurus* from the field, Eve Pieri and Isabel Manley for expert help with beetle husbandry, and the members of the Moczek and Newton Labs for constructive comments. This work was supported in part through generous funding from the National Science Foundation [Grant no. 2243725 and 1901680 to APM] and a National Science Foundation Graduate Research Fellowship [Grant No. 2141416 to JAJ]. The opinions, interpretations, conclusions, and recommendations are ours and are not necessarily endorsed by the National Science Foundation.

## Ethics disclosure statement

No IRB ethics approval was needed for the work presented in this study.

